# The neural correlates of semantic and grammatical encoding during sentence production in a second language: evidence from an fMRI study using syntactic priming

**DOI:** 10.1101/2021.04.11.438529

**Authors:** Eri Nakagawa, Takahiko Koike, Motofumi Sumiya, Koji Shimada, Kai Makita, Haruyo Yoshida, Hirokazu Yokokawa, Norihiro Sadato

**Author notes:** **Correspondence:** Norihiro Sadato, (NS).

## Abstract

Japanese English learners have difficulty speaking Double Object (DO; give B A) than Prepositional Object (PO; give A to B) structures which neural underpinning is unknown. In speaking, syntactic and phonological processing follow semantic encoding, conversion of non-verbal mental representation into a structure suitable for expression. To test whether DO difficulty lies in linguistic or prelinguistic process, we conducted functional magnetic resonance imaging. Thirty participants described cartoons using DO or PO, or simply named them. Greater reaction times and error rates indicated DO difficulty. DO compared with PO showed parieto-frontal activation including left inferior frontal gyrus, reflecting linguistic process. Psychological priming in PO produced immediately after DO and vice versa compared to after control, indicated shared process between PO and DO. Cross-structural neural repetition suppression was observed in occipito-parietal regions, overlapping the linguistic system in pre-SMA. Thus DO and PO share prelinguistic process, whereas linguistic process imposes overload in DO.

## 1. Introduction

Speaking is an automatic yet highly complex process. According to one widely cited model of speech production (Figure 1), it involves the generation of a preverbal message (*conceptualization*), translating it into a grammatical linguistic form (*formulation*), and articulating the phonetic plan (*articulation*) (Levelt, 1989). During conceptualization, *semantic encoding* occurs, which converts a non-verbal mental representation of the entity to be expressed (*reference*) into a semantic structure suitable for expression (*sense*) (Menenti et al., 2012a). Therefore, *sense* is the interface between *conceptualization* and *formulation*. Moreover, the *formulation* process involves grammatical encoding (Bock and Levelt, 1994), whereby syntax, the rules used to construct sentences (in specific languages) (Chomsky, 1957), is computed. Importantly, grammatical encoding is “no more accessible to conscious experience than the corresponding comprehension” (Bock and Levelt, 1994), and thus is a highly automatized process that may be linked to subconscious semantic encoding or the conceptualization process.

**Figure 1.**
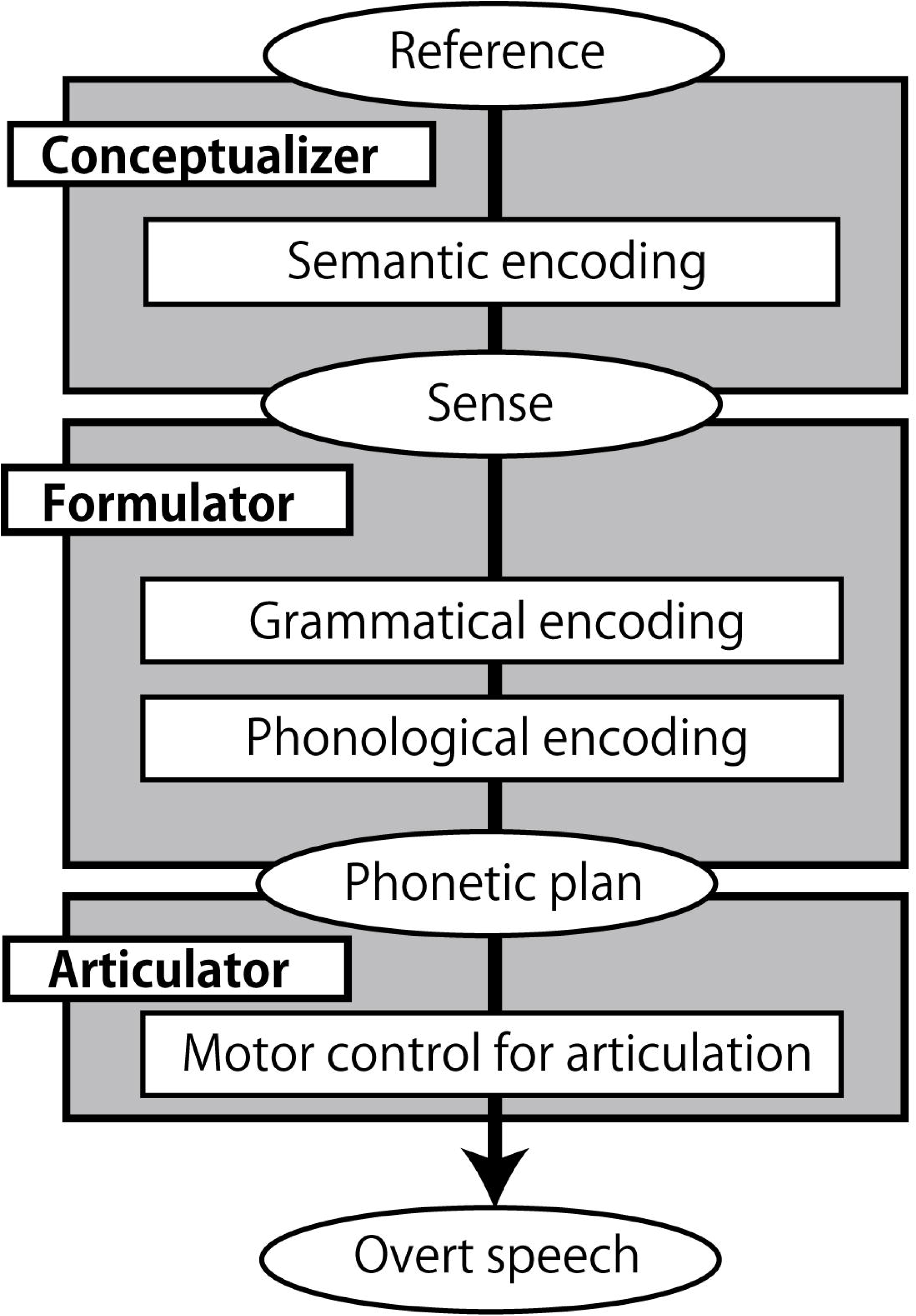
A speech production model. The model is based on (Levelt, 1989), (Bock and Levelt, 1994), and (Menenti et al., 2012a). Boxes represent processing components, and ellipses represent products and inputs of the processing components.

While speech production is automatic, speaking a second language (L2) is not as easy as speaking a first language (L1). Among the processing steps in speech production (Figure 1), it is not clear at what point the difficulty emerges in L2. Previous studies suggest that syntactic (grammatical) processing acts as a bottleneck for L2 learners. While L2 lexico-semantic processing gradually becomes native-like with higher proficiency (Hahne, 2001; Hahne and Friederici, 2001; Stein et al., 2009), reaching a native-like level for grammatical processing is difficult (Ullman, 2001; Clahsen and Felser, 2006c, 2006b). For example, unlike native English speakers, L2 learners do not show sensitivity to grammatical violations such as subject-verb number disagreements (e.g., *^1^The bridge to the *island were* about ten miles away) (Jiang, 2004). Additionally, for some aspects of grammar, neural processing becomes L1-like with higher proficiency (Ojima et al., 2005; Rossi et al., 2006), but complex syntactic structures (e.g., Which book did Mary think John believed the student had borrowed?; Clahsen & Felser, 2006c) are processed differently by L2 speakers (Marinis et al., 2005; Felser and Roberts, 2007).

L2 learners may under-use syntactic information while depending on lexical-semantic and pragmatic information, which leads to shallower and less detailed sentence processing (the shallow structure hypothesis) (Clahsen and Felser, 2006a, 2006b). Among the event-related potential (ERP) components related to grammatical processing, it has been shown that those that appear within an early time window, such as the early left anterior negativity (ELAN) or the left anterior negativity (LAN), are not seen in L2 learners (Kotz, 2009). This suggests that, unlike native speakers, L2 learners are not able to use syntactic information immediately. However, it is not yet clear why L2 speech production poses such a challenge, or how it is related to semantic and grammatical encoding or underlying neural mechanisms.

There is an interesting case in native Japanese speakers learning English as an L2. A two-character one-object scene depicting a “giving event” can be described with at least two structures using a ditransitive verb (e.g., *give*), including a Double Object (DO; e.g., He gives her the present) or Prepositional Object (PO; e.g., He gives the present to her) structure. Unlike native English speakers, Japanese English learners show a strong preference for PO over DO (Morishita, 2011, 2013; Morishita et al., 2011; Nakagawa et al., 2013), even though the essential semantic message underlying these two structures is identical. This suggests that Japanese English learners are processing PO and DO differently.

The neural underpinning of the difference in DO and PO processing is unknown. Specifically the similarity and difference of DO and PO processing along with the conceptualizer-formulator-articulator sequences have not been explored. To test if the difficulty of DO process resides in linguistic process or in prelinguistic semantic encoding process, we conducted functional MRI study with the sentence completion task. Each task trial was triggered by the cartoon explaining the situation thus providing the *reference*. Participants were required to utter the completed sentences in either DO or PO form. The trials were conducted serially, along with the control condition in which conceptualization and formulation components were eliminated. To depict the common neural processes between PO and DO, we utilized repetition suppression (Grill-Spector and Malach, 2001; Noppeney and Price, 2004; Grill-Spector et al., 2006; Auksztulewicz and Friston, 2016; Grotheer and Kovács, 2016; Larsson et al., 2016) across PO and DO, expecting a decrease in neural activity caused by repetitive exposure to the common properties beween PO and DO. The difference between the two structures was depicted by the classical subtraction method. We hypothesized that the difference is seen in the later process of the sentence production, that is, in the formulation process.

## 2. Materials and Methods

### 2.1. Participants

Thirty Japanese EFL learners, all college educated (16 female participants and 14 male participants; mean age ± standard deviation (SD) = 22.07 ± 2.78 years), participated in the experiment. All participants had normal or corrected-to-normal vision, normal hearing, and were right-handed according to the Edinburgh Handedness Inventory (Oldfield, 1971). Participants reported no history of neurological or psychiatric illness. The average age of English language acquisition (i.e., the age at which formal English language instruction was initiated) was 9.83 ± 4.14 years old. Ten participants reported that they had experience staying in an English-speaking environment for more than one month, with the duration of stay ranging from 2 to 122 months (mean ± SD = 13.93 ± 34.04 months). The Versant English Test (Pearson Education, Inc., 2011) was used to assess participants’ English proficiency. The Versant English Test is a 15-minute computerized exam that measures the user’s spoken English skills. See Table 1 for participants’ demographic information.

**Table 1.**
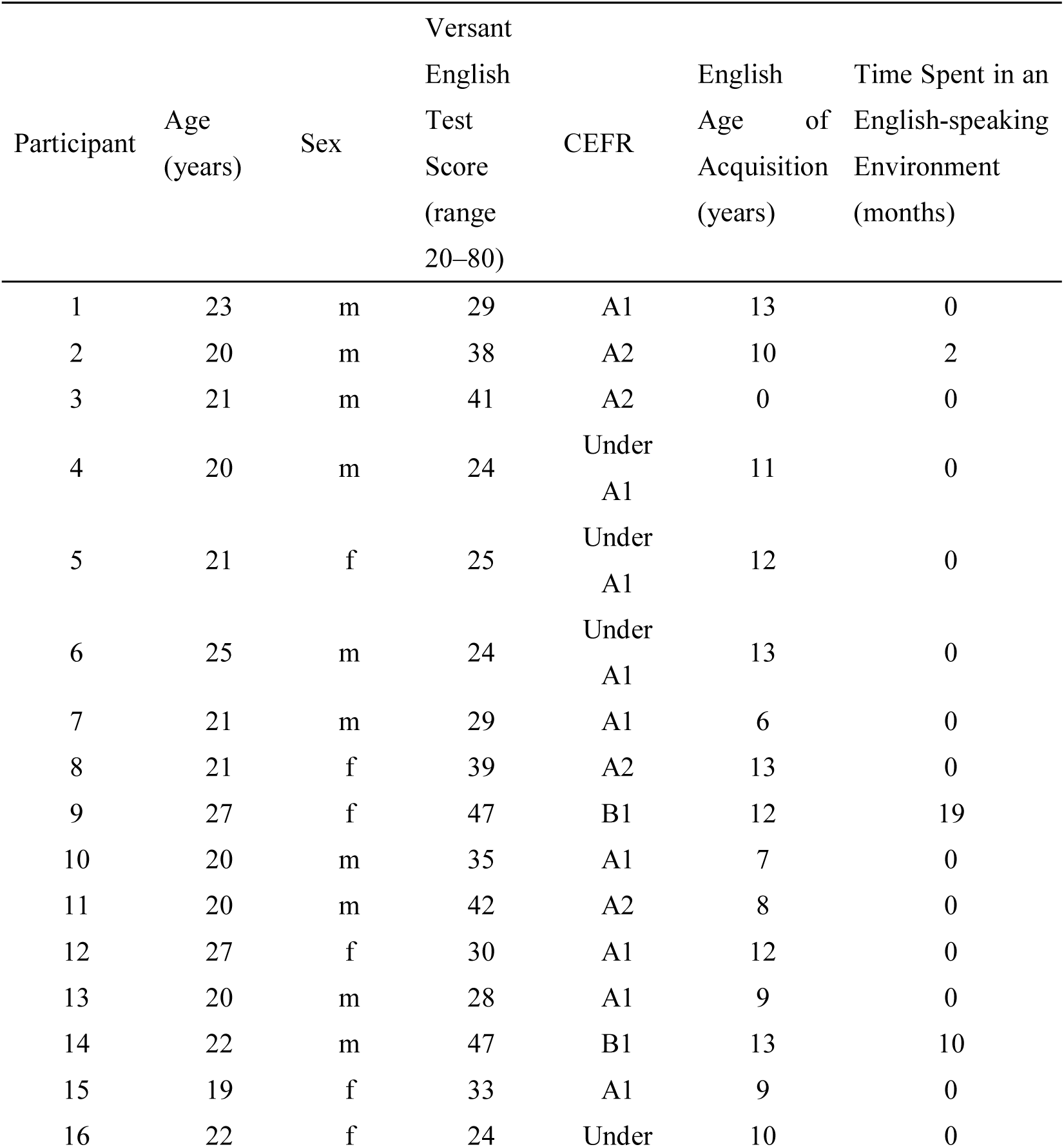
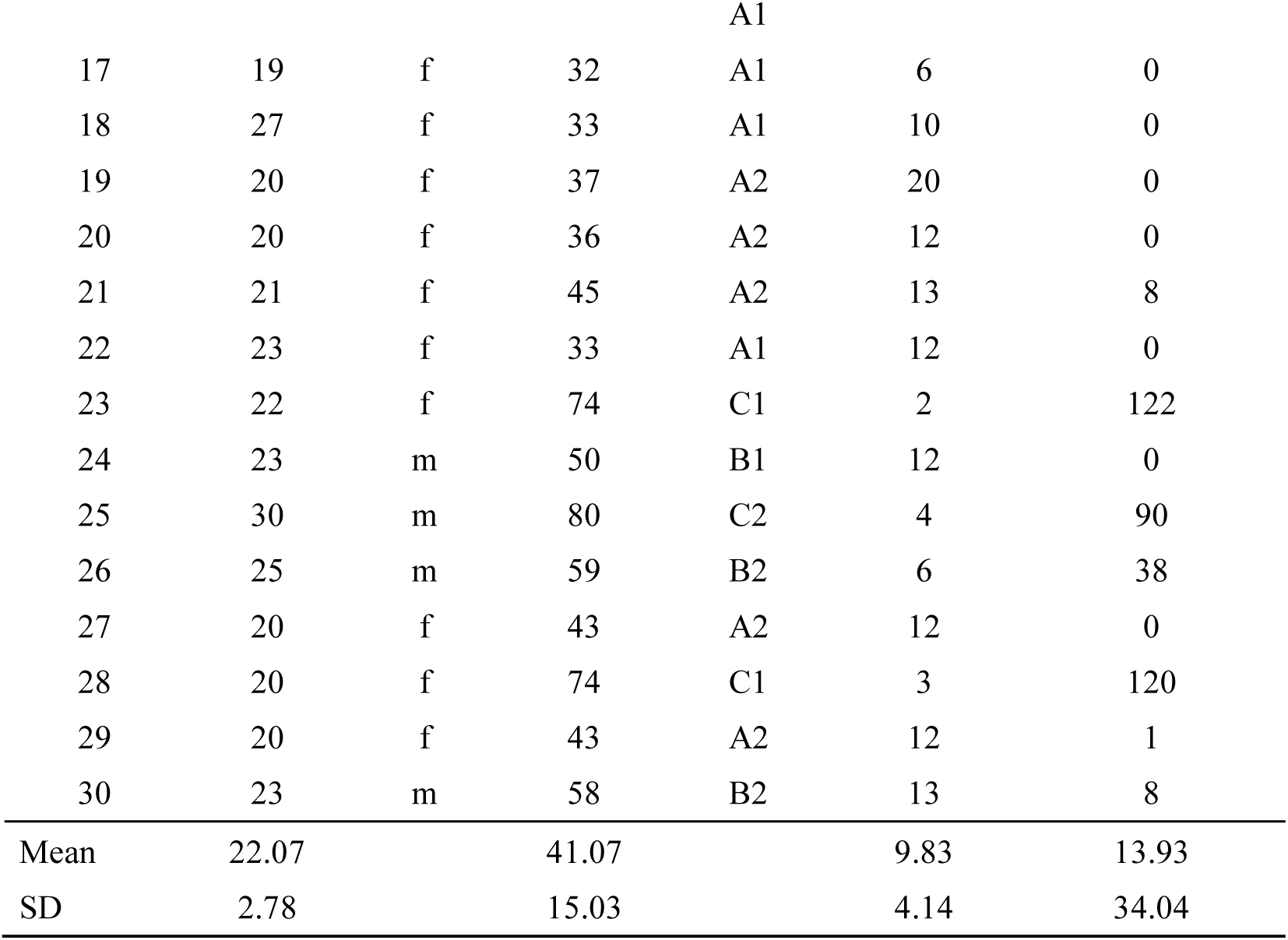
Demographic data of all participants. Note: The Common European Framework of Reference for languages (CEFR), is a guideline used to describe the achievement level of learners of foreign languages. It divides learners into six divisions as follows: A1, A2, B1, B2, C1, and C2, whereby A1 represents the least proficient and C2 indicates the most proficient. The CEFR scores were determined by the Versant English Test scores.

The protocol was approved by the Ethical Committee of the National Institute for Physiological Sciences, Japan. Experiments were undertaken in compliance with national legislation and the Code of Ethical Principles for Medical Research Involving Human Subjects of the World Medical Association (Declaration of Helsinki). All participants gave their written informed consent for participation.

### 2.2. Experimental design

We adopted an event-related design for the fMRI experiment. The trial order was pseudo-randomized to optimize the efficiency of the design (Dale, 1999; Friston et al., 1999). There were six runs in total and each run included 48 trials. The total number of trials throughout the experiment was 288. One run consisted of four blocks of 12 consecutive 6000-ms trials that required an oral response. The four blocks were separated by two consecutive 6000-ms rest trials. An 18-s and 12-s baseline epoch were conducted before the first trial and after the last trial, respectively. Each run lasted for approximately 6 minutes (354 s).

Each target item served as the prime sentence for the next target item (the running priming paradigm, Menenti, Gierhan, Segaert, & Hagoort, 2011; Menenti, Petersson, et al., 2012; Menenti, Segaert, & Hagoort, 2012; Segaert, Kempen, Petersson, & Hagoort, 2013). Figure 2 shows an example of a trial sequence. “P”, “D”, and “N” indicate PO, DO, and No Structure trials, respectively. “R” indicates Rest trials, in which a cross mark on a black screen was presented for 6000-ms. The target trial (the “present” utterance) is notated in upper-case font, with its preceding trial (the utterance in the previous trial) in lower-case font. For instance, when the target trial had a DO structure and was preceded by a PO structure trial, the designation would be pD. As we are interested in the effect of the preceding trial on the present trial (i.e., the priming effect), a 3 x 3 design with the factors of present utterance (P, D, or N) and previous utterance (p, d, or n) was used.

**Figure 2.**
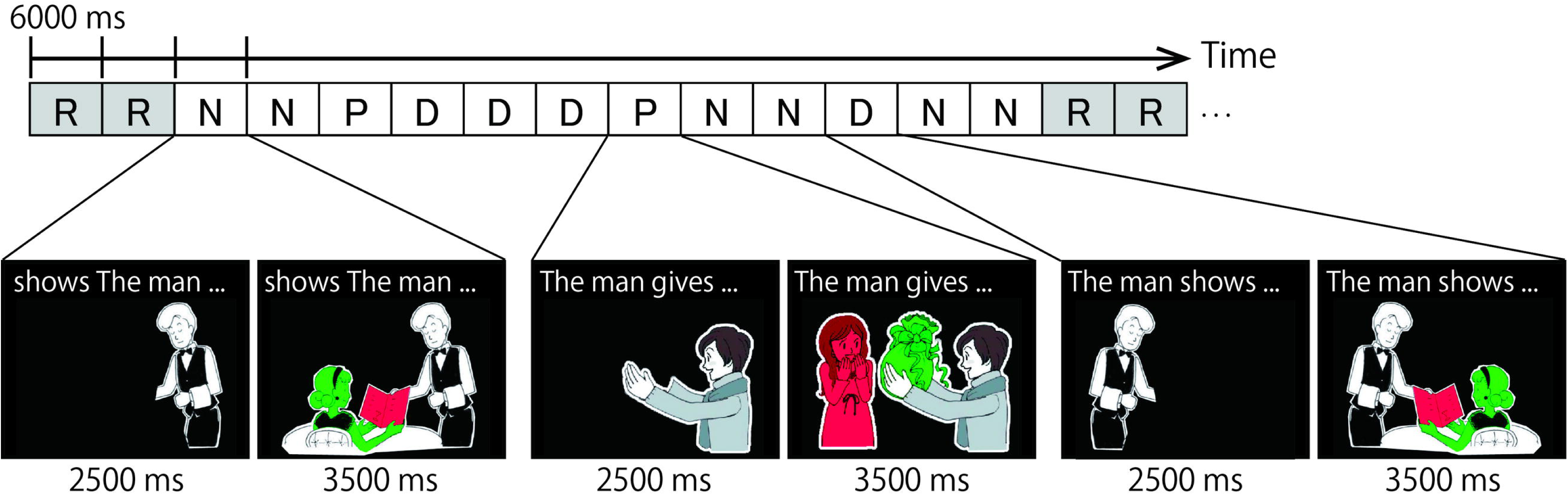
Example of a task sequence. R, N, P, and D represent Rest, No structure, PO, and DO conditions, respectively. Each condition was presented for 6 s and consisted of two parts. In the first part, a cartoon of the actor was presented with a sentence fragment printed in the upper part of the screen. In the second part, participants were asked to describe the colored pictures by referring to the green object before the red object.

### 2.3. Task and procedure

The task was to describe a cartoon by completing the sentence fragment printed above it. Each trial had a duration of 6000 ms and consisted of two parts. The first part required participants to silently read the sentence fragment (e.g., The man gives…) within 2500 ms. The sentence fragments shown above the cartoons were in a specific grammatical order, and the picture depicted the agent of the action only. After 2500 ms, the colored objects, which represented the theme object and the recipient of the action were presented for 3500 ms. In this second part of the trial, participants overtly described the cartoon by completing the sentence fragment they had read silently. The two newly presented objects were colored either green or red, and participants were instructed to describe the scene by referring to the green object before the red object (the stoplight paradigm; Menenti et al., 2011; Menenti, Petersson, et al., 2012; Menenti, Segaert, et al., 2012; Schoot, Menenti, Hagoort, & Segaert, 2014; Segaert et al., 2013; Segaert, Menenti, Weber, & Hagoort, 2011; Segaert, Menenti, Weber, Petersson, & Hagoort, 2012; Segaert, Wheeldon, & Hagoort, 2016). This manipulation determined which sentence structure (PO or DO) was produced. For example, when “the present” was shown in green and “a girl” in red, the correct spoken response would be “the present to the girl”, which is a PO response. In some trials, ungrammatical fragments (e.g., gives the man …) were presented above the picture. In these No Structure trials, participants were instructed to simply name the two objects by referring to the green one before the red one (e.g., “present, girl”). Participants were instructed to fixate on a cross that appeared in the center of the screen after every 12 trials (Rest).

### 2.4. Stimuli

#### 2.4.1. Selection of materials

Three dative verbs (*give, sell,* and *show*) were selected from an English vocabulary familiarity database, based on the rating of 810 Japanese college students learning EFL (Yokokawa, 2006). Vocabulary familiarity refers to how often people (Japanese English learneres) perceive that they hear or see a word. In contrast to the actual frequency of a word in newspapers, magazines, or the language corpus as a whole, familiarity reflects internal or mental frequency, and is scored between 1 (least familiar) and 7 (most familiar). The mean familiarity scores for all three dative verbs used in the present study were as follows: give = 6.69, sell = 5.61, and show = 6.44. By definition, dative verbs must be followed by two objects, including the entity that is acted upon (theme) and the entity that is the beneficiary of the action. Dative verbs are used to construct PO and DO structures and cannot be used in a simple transitive form (*\*He gives her*). Cartoons that depicted a ditransitive event involving two actors and one object were used as stimuli. For each of the three dative verbs, four cartoons were selected from a website that supplied materials free of charge (https://minnanokyozai.jp/). The cartoons were all describable using the dative verbs in both PO and DO phrases. All but one cartoons were identical to those used in our previous study (Nakagawa et al., 2013). Using Adobe Photoshop CS5 (Version 12.0×64; Adobe Systems Incorporated, San Jose, CA, USA), the recipient and the theme object of the action in the cartoons were colored either green or red. We created mirrored versions for all of the cartoons. There were 24 different cartoon stimuli in total.

#### 2.4.2. Stimulus presentation

Presentation software (Version 19.0, Neurobehavioral Systems, Albany, CA) was used to present the visual stimuli. A liquid crystal display (LCD) projector (CP-SX12000J; Hitachi, Tokyo, Japan) located outside the scanner room projected the stimuli through a waveguide onto a half-transparent screen behind the scanner, which the participants viewed via a mirror attached to the head coil of the scanner. The spatial resolution of the projector was 1024 × 768 pixels, with a 60 Hz refresh rate. While the exact value varied depending on the subject’s head size, the screen was approximately 190 cm from the participants’ eyes. The visual angle of stimulus size was 13.06° × 10.45°. Spoken responses were collected via a MR-compatible, noise-cancelling optical microphone system (LITEMIC™ 3140 [FOMRI-II]; Optoacoustics, Israel) attached close to the mouth.

### 2.5. Experimental procedure

Participants were informed that the purpose of the experiment was to examine how fast Japanese EFL learners could describe a given picture in English. After providing informed consent, participants underwent a training session outside of the MR scanner. The training session consisted of three parts. The objective of the training was to familiarize the participants with the objects they would have to name to facilitate the speed of word retrieval during the experiment. This was essential because participants were all Japanese EFL learners, and we were concerned that the picture description may not have been completed within 3500 ms if word retrieval was too slow.

In the first part of the training session, participants named single objects extracted from the target cartoons. Cartoons appeared one at a time at a rate of one every 2 s. The cartoons for the training trials were not colored red or green. Example words were printed below the cartoons. Participants could simply read them out loud or name the cartoons in any way they liked. They underwent another training run without any words printed below the cartoons.

In the second part of the training session, participants viewed the 24 cartoons individually as per the actual experiment. They were instructed to describe the cartoons by silently reading the sentence fragment above the cartoon, and then saying the remaining part of the sentence out loud. The purpose of this part of the training session was to familiarize participants with the stoplight paradigm, i.e., referring to the green object before the red object. During this part of the training, the experimenter presented the pictures individually without time constraints. Participants received feedback (correct/incorrect PO and DO construction) from the experimenter after each practice trial.

In the third part of training, participants underwent a practice run of 12 trials. We used stimuli from the actual experiment, but using a different trial order. This training was performed once outside and once inside the MRI scanner. After completing the training session, all participants were able to respond to the stimuli within 3500 ms.

#### 2.5.1. MRI data acquisition

A 3 Tesla (3T) whole-body scanner (Verio; Siemens Erlangen, Germany) with a 32-element phased-array head coil was used to acquire fMRI and anatomical data. To obtain T2*-weighted (functional) images, we employed a multi-band echo-planar imaging (EPI) sequence that collected multiple EPI slices simultaneously and reduced the volume repetition time (TR) (Moeller et al., 2010). We utilized the following sequences to cover the whole brain: TR = 500 ms, acquisition time (TA) = 500 ms, echo time (TE) = 30 ms, flip angle (FA) = 90°, field of view (FOV) = 192 mm, 64 × 64 matrix; voxel dimensions = 3.0 × 3.0 × 3.0 mm with a 0.5 mm gap, number of slices = 42; and multiband factor = 6. Anatomical images were acquired using a T1-weighted Magnetization-Prepared Rapid-Acquisition Gradient Echo (MPRAGE) sequence with the following parameters: TR = 2400 ms, TE = 2.24 ms, FA = 8°, FOV = 256 mm, and voxel dimension = 0.8 × 0.8 × 0.8 mm, number of slices = 208.

### 2.6. Data processing

#### 2.6.1. Behavioral data analysis

The spoken responses were transcribed and coded for errors. Responses were scored as PO if the subject and verb were followed by a noun phrase that behaved as a patient/theme, and then by a prepositional phrase beginning with *to*, which behaved as the beneficiary. It was scored as DO if the verb in the utterance was immediately followed by a noun phrase that behaved as the beneficiary, and then by a noun phrase, which behaved as the patient/theme. Responses without a determiner (such as *the* or *a*) in the PO or DO trials were scored as errors. Responses were also considered errors when the colored objects were not referred to in the correct order (i.e., green before red), utterances had one object missing (e.g., The man is giving a present), utterances had an incorrect word order (e.g., The man give to a girl a present), or a verb agreement error. Based on these criteria, we calculated the error rate for each experimental condition.

Reaction time (RT) was defined as the speech production latency following the onset of the presentation of the colored objects. A beep that was time locked to the onset of each trial was simultaneously recorded with the participants’ responses. The beep was used as a marker for analysis and was not presented to the participants. RTs were only calculated for the trials with correct responses.

#### 2.6.2. Statistics

The general linear model (GLM) repeated measures in IBM SPSS Statistics for Windows (Version 20.0. IBM Corp, Armonk, NY) was used to analyze error rate and RT data. Results of analyses were considered statistically significant if *p* < .05. Figures (bar graphs) were created using GraphPad Prism (Version 5.03) for Windows (GraphPad Software, San Diego California USA; www.graphpad.com).

#### 2.6.3. fMRI data analysis

Image processing and statistical analyses were performed using the Statistical Parametric Mapping package (SPM12; Wellcome Trust Centre for Neuroimaging, London, UK) implemented in MATLAB (R2016b, MathWorks, Natick, MA, USA). A total of 708 volumes were acquired per run. The first 12 out of 708 volumes of each run were discarded to allow for stabilization of the magnetization. The remaining 696 volumes per run were used for analysis. All volumes were realigned for motion correction. The anatomical image was co-registered to the mean image of the functional images. The co-registered anatomical image was normalized to the Montréal Neurological Institute (MNI) T1 image template (ICBM152) (Evans et al., 1993; Friston et al., 1995) using a nonlinear basis function. The same normalization parameters were applied to all of the EPI volumes.

As head motion is known to affect fMRI results, we conducted rigid artifact removal with FSL’s FIX tool (FMRIB’s ICA-based Xnoiseifier) (Griffanti et al., 2014; Salimi-Khorshidi et al., 2014). In this study, we modified the MultiRunFIX developed by the HCP Human Connectome Project (HCP) pipeline (Glasser et al., 2018) so that it could be appled to data preprocessed by the SPM software to remove structured noise. The independent components were extracted by Multi-Run sICA (spatial independent component analysis) from the normalized EPI data using a nonlinear basis function which were concatenated from six runs. Linking of the 6-run data was performed since it is more advantageous to have more time points in order to improve the noise and signal separation performance by spatial ICA. This reduces the risk of removing not only noise but also task-related activities due to low separation performance. To the extracted independent components, automatic labeling based on machine learning was not performed, but we performed hand classification of the ICA components (Griffanti et al., 2017). The concatenated data were divided and returned to the data of each run, and data analysis after this point was done using SPM. The images were spatially smoothed with an 8-mm full-width at half-maximum Gaussian kernel along the x, y, and z axes.

Statistical analysis of the functional imaging data was conducted in two steps. At the first level, single subject task-related activation was analyzed using a GLM (Friston et al., 1994; Worsley and Friston, 1995). Nine regressors of interest and one regressor of no interest were included in the design matrix for each individual subject. The regressors of interest modeled the experimental conditions. Depending on the previous trial, there were three conditions each for PO (pP, dP, nP), DO (pD, dD, nD), and No Structure responses (pN, dN, nN). The onset of these regressors were specified at the beginning of the second trial cue with 0 duration. The regressor of no interest was added to model out the utterance related effect. The onset of this regressor was specified at the voice onset with 0 duration. Whenever there was no response, the onset was set at the end of the trial, which was 3.5s after the second trial cue.

The weighted sum of the parameters estimated in the first-level analysis consisted of “contrast” images that were used for the random effects group analysis (Friston et al., 1996). In this second-level analysis, we used a factorial design (within-subjects one-way analysis of variance [ANOVA]) with nine contrast images (pP, dP, nP, dD, pD, nD, pN, dN, nN) from each participant. The threshold for significance of the SPM{t} was set at *p* < .05 with a family-wise error (FWE) correction at the cluster level for the entire brain with an uncorrected height threshold of *p* < .001 (Friston et al., 1996). We evaluated the following two contrasts: First, to reveal the neural substrates related to the prelinguistic process (semantic encoding), the contrast of repetition suppression of cross-structural priming with the effect of N priming removed for PO ([dP < nP] + [nN < dN]) and DO ([pD < nD] + [nN < pN]) was evaluated using a conjunction null analysis (Friston et al., 2005). The regions showing suppression in both PO to DO and DO to PO conditions are related to semantic encoding as they reflect a common processing between PO and DO. Second, to reveal the additional cognitive load for DO compared to PO production, we evaluated the difference between PO and DO (nD > nP). We hypothesized that this contrast would reflect regions related to grammatical encoding. Brain regions were anatomically defined and labeled according to a probabilistic atlas, Anatomy Toolbox ver 3.0 (Eickhoff et al., 2005, 2006, 2007). The activation patterns were rendered on the skull-stripped ICBM152 MNI template pre-installed in the SPM package using MRIcroN (Version 1). We evaluated brain activation after excluding any activation outside the gray matter with the masking procedure.

### 2.7. Methodological considerations

The mirrored versions were created because the composition of the objects in the stimuli may have caused bias. For example, if the actor was always depicted on the right, the theme object in the middle, and the recipient on the left, participants may have formed an actor-theme-recipient structure, i.e., PO structure by scanning the cartoon from right to left and naming them in order. The position of the theme object was almost always in between the actor and the recipient; nevertheless, the positions of the actor and recipient were controlled by using both original and mirrored versions of all stimuli. A less biased stimuli that eye scanning preferences do not potentially prime a word order could be created by depicting three objects in a random position (Kootstra and Doedens, 2016). This was not adopted because in such case speakers must create the concept of the cartoon (i.e., conceptualization, Bock & Levelt, 1994; Levelt, 1989) by themselves and thus it would affect the following semantic and grammatical encoding processes, especially for those with low proficiency. It was important that participants finished describing the cartoons within 3.5 seconds due to our experimental design. Thus, we prioritized the cartoons to depict a natural dative scene instead of arranging the objects randomly.

## 3. Results

### 3.1. Behavioral results

#### 3.1.1. Correcting for inter-subject variability

There were four covariates, as follows: The Versant English test score (proficiency), age, age of acquisition, and the amount of time spent in an English-speaking environment (i.e., amount of exposure), that were considered to account for the differences in inter-subject variability. We conducted an a priori test to examine if these four covariates met the assumptions for analysis of covariance (ANCOVA). We found that none of the four covariates explained the error rate data. For RTs, we found that the Versant English test score significantly explained the data (*p* = .026), while the other three covariates did not. As this suggested that the Versant English test score was not independent from the RT data, we conducted a 3 x 2 ANCOVA with one covariate (Versant English test score) for RT data, and a 3 x 2 repeated measures ANOVA for the error rate data to investigate how the previous trial affected the present trial on target syntactic structures. The factors were the structure uttered in the previous trial, p, d, or n, and the structure uttered in the present trial, P or D. For these 3 x 2 conditions, we calculated the relative change from the No structure condition for both RT and error rate before conducting the ANCOVA and ANOVA.

#### 3.1.2. Error rate

Mauchly’s test indicated that the assumption of sphericity had been violated, and degrees of freedom were therefore corrected using Huynh-Feldt estimates of sphericity for the “previous structure” factor (Chi-Square (2) = 8.17, *p* = .017). There were significant main effects of the structure type in the present utterance (*F*(1, 29) = 11.384, *p* = .002) and the previous utterance (*F*(1.674, 48.554) = 5.789, *p* = .008) on error rate. There was a trend for an interaction between the present and previous utterance on error rate (*F*(2, 58) = 3.084, *p* = .053). These results indicate that, irrespective of the previous utterance, error rate was higher for DO than PO trials. Also, irrespective of the present utterance (*p* = .002), error rate was higher when the previous utterance was a non-syntactic response than when it was a syntactic response (p < n, *p* = .002; d < n, *p* = .025) (Figure 3a). In other words, there was a facilitatory effect when syntactic structures (irrespective of same or different structures) were repeatedly produced compared to when it was produced after a non-syntactic structure, that is, a syntactic priming effect for pP and dD trial pairs, and a cross-structural priming effect for trial pairs involving a PO-DO or a DO-transition.

**Figure 3.**
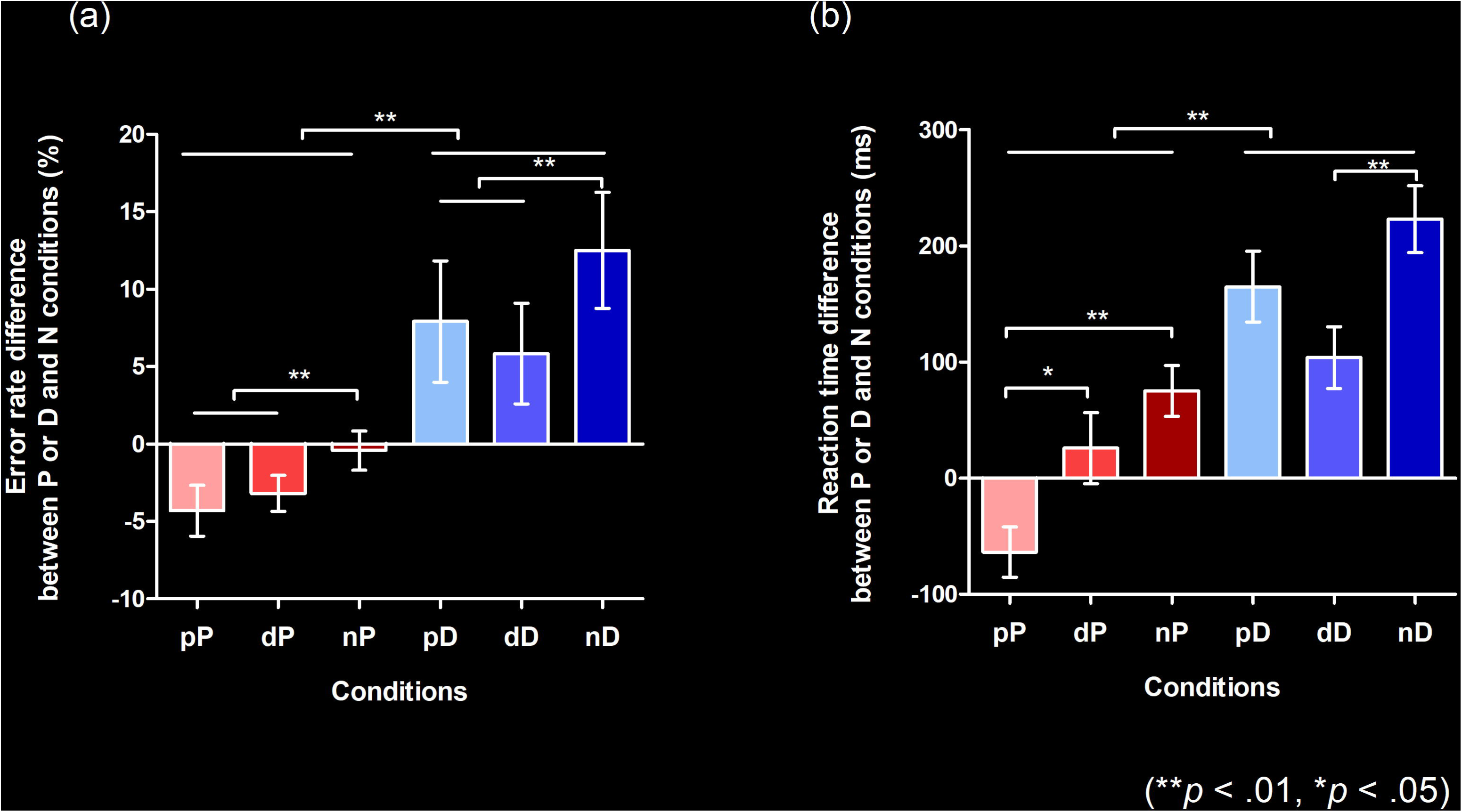
Behavioral results (\*\**p* < .01, \**p* < .05; Bonferroni-corrected). Both error rate (a) and reaction time (b) data in this figure show the relative difference compared to the No structure condition. (a) The relative error rate from the N condition for P and D conditions. Error bars indicate the standard error of the mean (SEM). All responses other than PO (verb followed by a noun phrase that behaved as a patient/theme + a prepositional phrase that behaved as the beneficiary) or DO (verb immediately followed by a noun phrase that behaved as the beneficiary + a noun phrase that behaved as the patient/theme) were coded as errors. (b) The relative reaction times from the N condition for P and D conditions. Error bars indicate the SEM. Reaction time indicates the time measured from the point when the colored objects were presented to when participants started uttering the first word. Error responses were excluded from the data used in this figure. P, D, and N represent Prepositional Object trials (PO), Double Object trials (DO), and No structure trials, respectively. The target trial is notated in upper case font, with the preceding trial (prime) in lower case font.

#### 3.1.3. Reaction Time (RT)

Mauchly’s test indicated that the assumption of sphericity had been met. The ANCOVA revealed a significant main effect of structure type in the present utterance on RT (*F*(1, 28) = 10.485, *p* = .003), and a significant interaction between structure type in the present and previous utterances on RT (*F*(2, 56) = 3.369, *p* = .042). There was a trend towards a main effect of structure type in the previous utterance on RT (*F*(2, 56) = 2.925, *p* = .062). Post hoc pairwise comparisons with Bonferroni’s correction revealed that RT in the PO condition when the previous trial was also PO was shorter than when the previous trial was DO (pP < dP, *p* = .036) or when the previous trial was No structure (pP < nP, *p* < .001). RT in the DO condition was shorter when the previous trial was also DO compared to when it was No structure (dD < nD, *p* < .001) (Figure 3b), but RT reduction was not seen when the previous trial was PO (dD < pD was not significant). This result shows that there was no cross-structural priming effect for RTs. The RT in the nN condition was significantly faster than nP and nD conditions (both *p* < .05), which indicates that participants were not automatically running grammatical processing in the control condition.

### 3.2. fMRI results

#### 3.2.1. Correcting for inter-subject variability

A priori test of the behavior data (RT), showed that only the Versant English Test score significantly explain the data. Based on this finding, we performed fMRI data analysis including the Versant English Test score as a covariate.

#### 3.2.2. Repetition suppression of syntax processing

The behavioral results for error rate showed a facilitatory effect of syntax processing due to repetition (i.e., pP < nP and dD < nD were significant). To identify the brain regions underlying this priming effect, we evaluated the repetition suppression of cross-structural priming contrast for PO ([dP < nP] + [nN < dN]) and DO ([pD < nD] + [nN < pN]), and using a conjunction analysis. Consequently, activation was observed in the fronto-parieto-occipital regions, including the pre-supplementary motor area (SMA), bilateral superior parietal lobule (SPL), and the bilateral inferior occipital gyrus (IOG) (Figure 4, Table 2).

**Figure 4.**
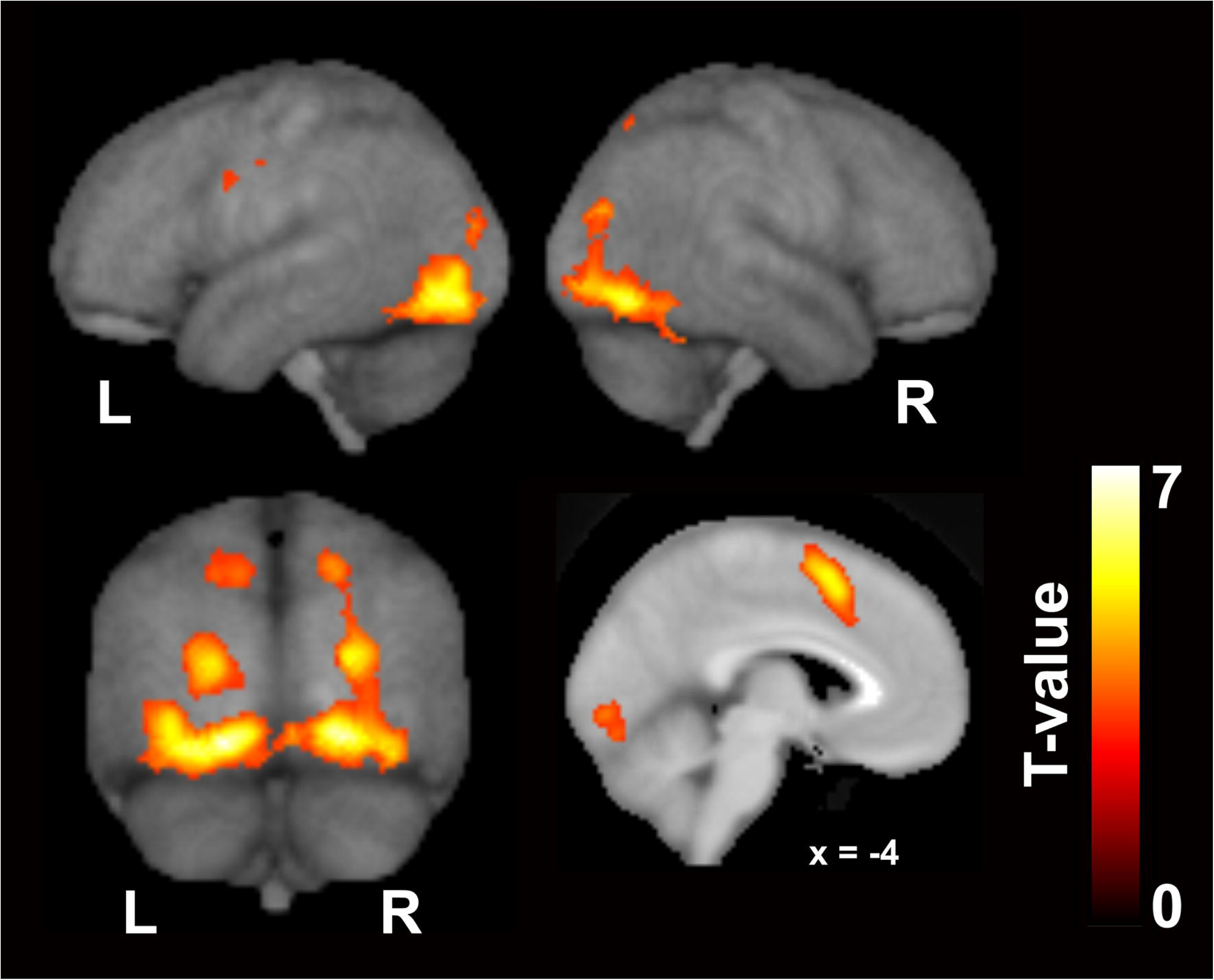
The regions showing repetition suppression by cross-structural priming, excluding the effect of N priming (conjunction null analysis of [dP < nP] + [nN < dN] and [pD < nD] + [nN < pN]). In this cross-structural priming contrast, we aim to investigate the priming effect due to repetition of the function of semantic encoding. FWE-corrected *p* < .05 at the cluster level, with a height threshold of *p* < .001, uncorrected. These regions reflect a common processing between PO and DO. The activation is rendered on the skull-stripped ICBM152 MNI template pre-installed in the SPM package.

**Table 2.**
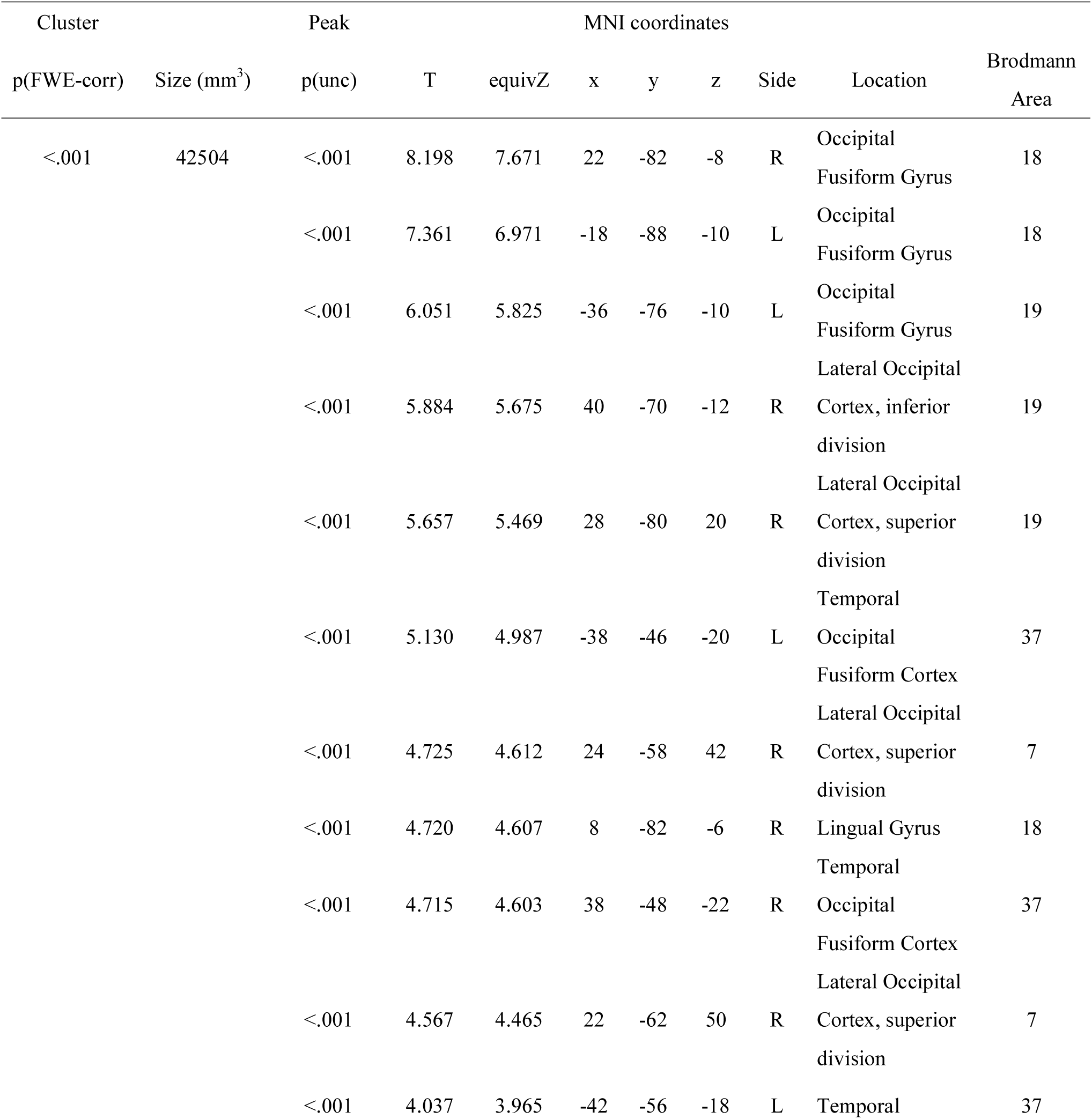
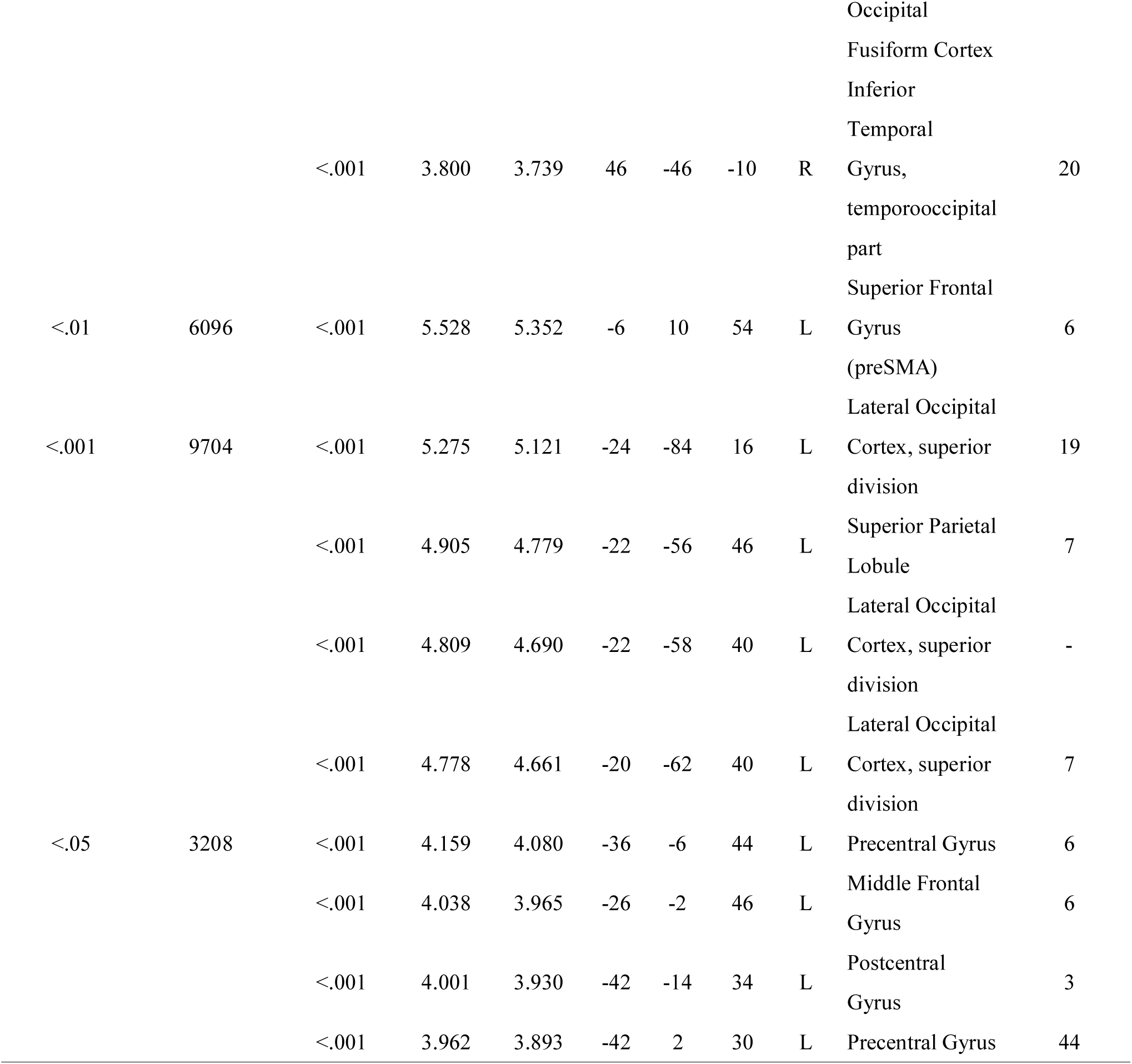
The regions showing repetition suppression of cross-structural priming, excluding the effect of N priming (conjunction of [dP < nP] + [nN < dN] and [pD < nD] + [nN < pN]).

#### 3.2.3. Difference between DO and PO production

The behavioral data demonstrated a clear difference between PO and DO production, whereby DO production was more demanding. To identify the corresponding regions involved, we compared the activation patterns for the P and D conditions (nD > nP). Activation was observed in the pre-SMA, bilateral inferior frontal regions, including the pars opercularis (BA44), particularly in the left hemisphere, and areas along the inferior frontal sulcus extending to the frontal pole. (Figure 5, Table 3).

**Figure 5.**
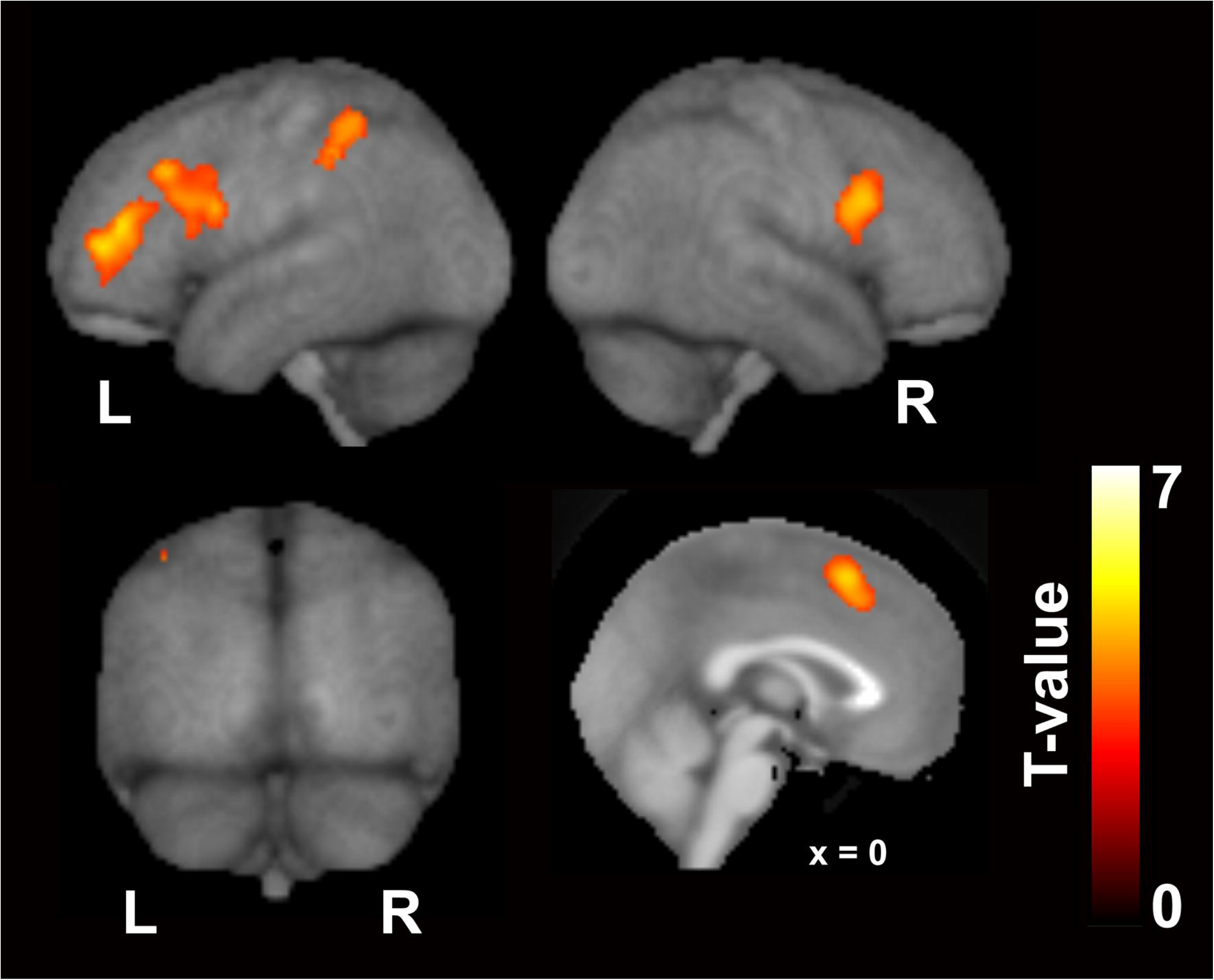
Regions showing greater activation in DO than in PO (nD > nP). Activation in these regions reflect a greater processing load for grammatical encoding in DO compared to PO production. FWE-corrected *p* < .05 at the cluster level, with a height threshold of *p* < .001, uncorrected. The activation is rendered on the skull-stripped ICBM152 MNI template pre-installed in the SPM package.

**Table 3.**
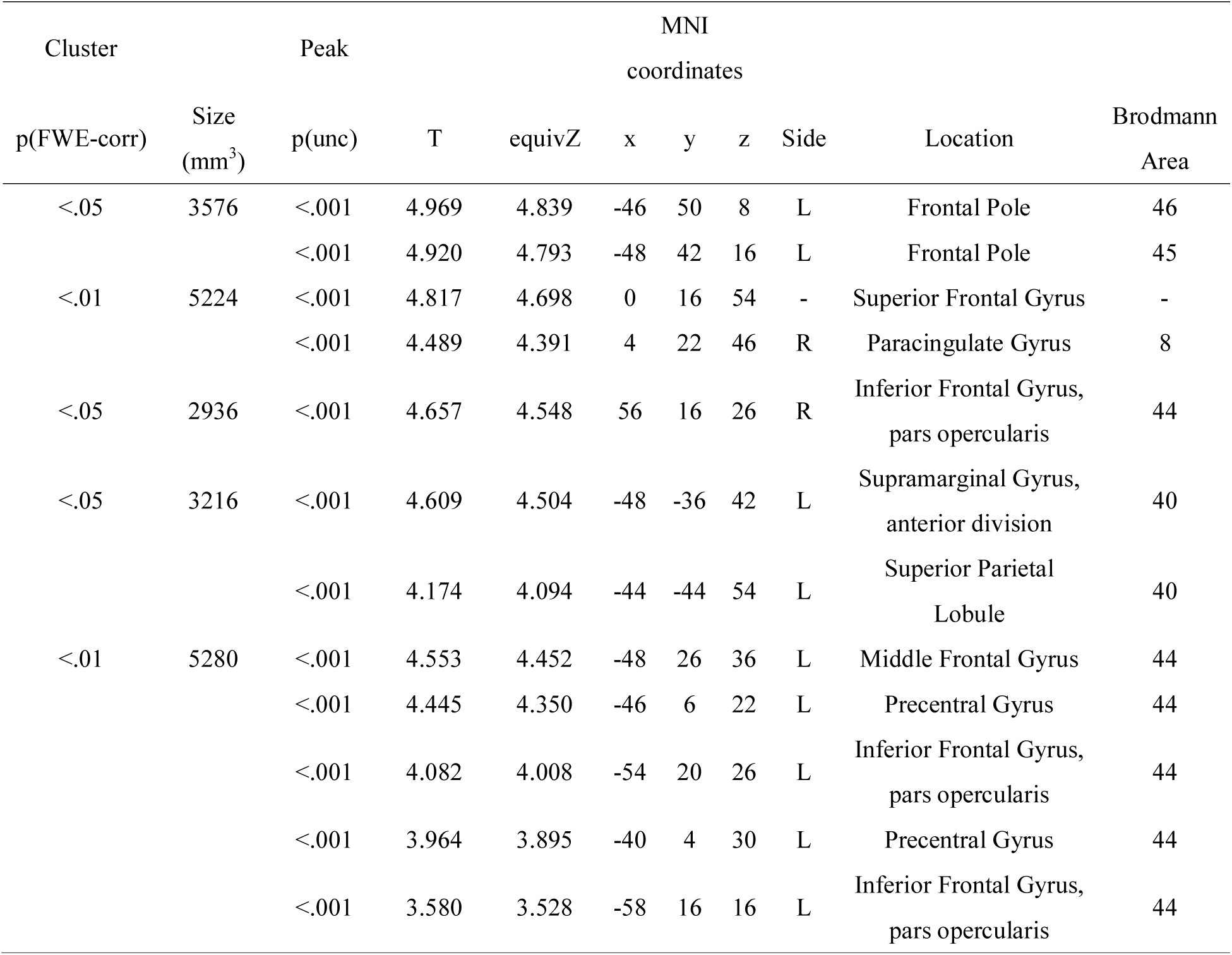
The regions showing greater activation in DO compared to PO (nD > nP).

## 4. Discussion

### 4.1. Neural substrates of L2 sentence production common to DO and PO

To depict the common neural substrates for DO and PO processing we utilized repetition suppression, which is a decrease in neural activity caused by repetitive exposure to the same properties (Grill-Spector and Malach, 2001; Noppeney and Price, 2004; Grill-Spector et al., 2006; Auksztulewicz and Friston, 2016; Grotheer and Kovács, 2016; Larsson et al., 2016). A previous fMRI study using repetition suppression reported that a widespread network of areas associated with language processing, including the left middle frontal gyrus, bilateral superior parietal lobes, and bilateral posterior temporal gyri, are related to semantic encoding, or, in other words, the construction of non-verbal mental representations of the sentence meaning (Menenti et al., 2012a). Conjunction analysis of the cross-structural repetition suppression showed activation in the bilateral IOG, bilateral SPL, and the pre-SMA, the last of which was also activated by DO-PO (nD > nP) comparison. This distribution of common contribution suggests these areas are responsible for the prelinguistic conceptualization process.

#### 4.1.1. IOG

The task in the present study was to describe a cartoon. The first step was to perceive and interpret the concept of the cartoon and specify the relational semantic structure, known as the thematic roles. Activation in the IOG likely reflects the visual perception of the stimuli. The bilateral occipital pole plays an important role in the identification of animated entities and the dynamic relationships between them (Morito et al., 2009). The present study used cartoons showing two characters dynamically interacting with each other, and thus it is reasonable that these areas were activated. In summary, these areas are related to the creation of the *reference* (the mental representation an utterance refers to) (Menenti et al., 2012a).

#### 4.1.2. SPL

Studies with L1 speakers of Dutch that used a picture description task with the stop light paradigm, which is the same task used in the present study, reported that the left SPL, bilateral middle temporal gyrus (MTG), and precuneus demonstrated repetition suppression when semantics were repeated (Menenti et al., 2012b). Menenti and colleagues focused on semantic encoding in particular, and reported that the bilateral SPL, precentral gyrus, left IFG, and posterior MTG exhibited repetition suppression effects for both *reference* and *sense* (the linguistic structure that interfaces meaning with linguistic form) (Menenti et al., 2012a). The bilateral SPL is involved in linguistic inference (Nieuwland et al., 2007; Monti et al., 2009). Menenti and colleagues hypothesized that repetition suppression in the SPL represents decreased requirement for inferences when *sense*, *reference,* or both are repeated. Similar to L1 speakers, semantic encoding in L2 recruited occipito-parietal regions. We speculate that the relationship between the characters (*reference* of the scene) is processed in the occipital areas while the creation of *sense* engages broader areas, including parietal areas.

#### 4.1.3. Pre-SMA

The output of semantic encoding (*sense*) is the input for the next step in speaking, which is grammatical encoding. In the case of L1 speaking, semantic, lexical, and syntactic processes involve partly overlapping but distinct brain networks (Menenti et al., 2012b). The pre-SMA is involved in both semantic encoding and grammatical encoding, the latter of which is DO dominant. The previous L1 study has shown that the pre-SMA is a semantic encoding-related area (cf. Menenti, Petersson, et al., 2012). Previous work has reported that this region is also involved in syntax-related tasks (Menenti et al., 2011, 2012b). For example, an L1 study investigating the comprehension of differentially complex syntactic structures reported that the left dorsal premotor cortex and left SMA were sensitive to syntactic complexity when a sentence included two animate characters whose semantic roles could be reversed (syntactic complexity due to reversibility) (Meltzer et al., 2010). These syntax-related areas were included in the regions representing semantic encoding in the present study, as well as in L1 studies (Menenti et al., 2012a, 2012b). We propose that the neural substrates underlying semantic and grammatical processes are partially overlapping in the pre-SMA of L2 learners (Figure 6).

**Figure 6.**
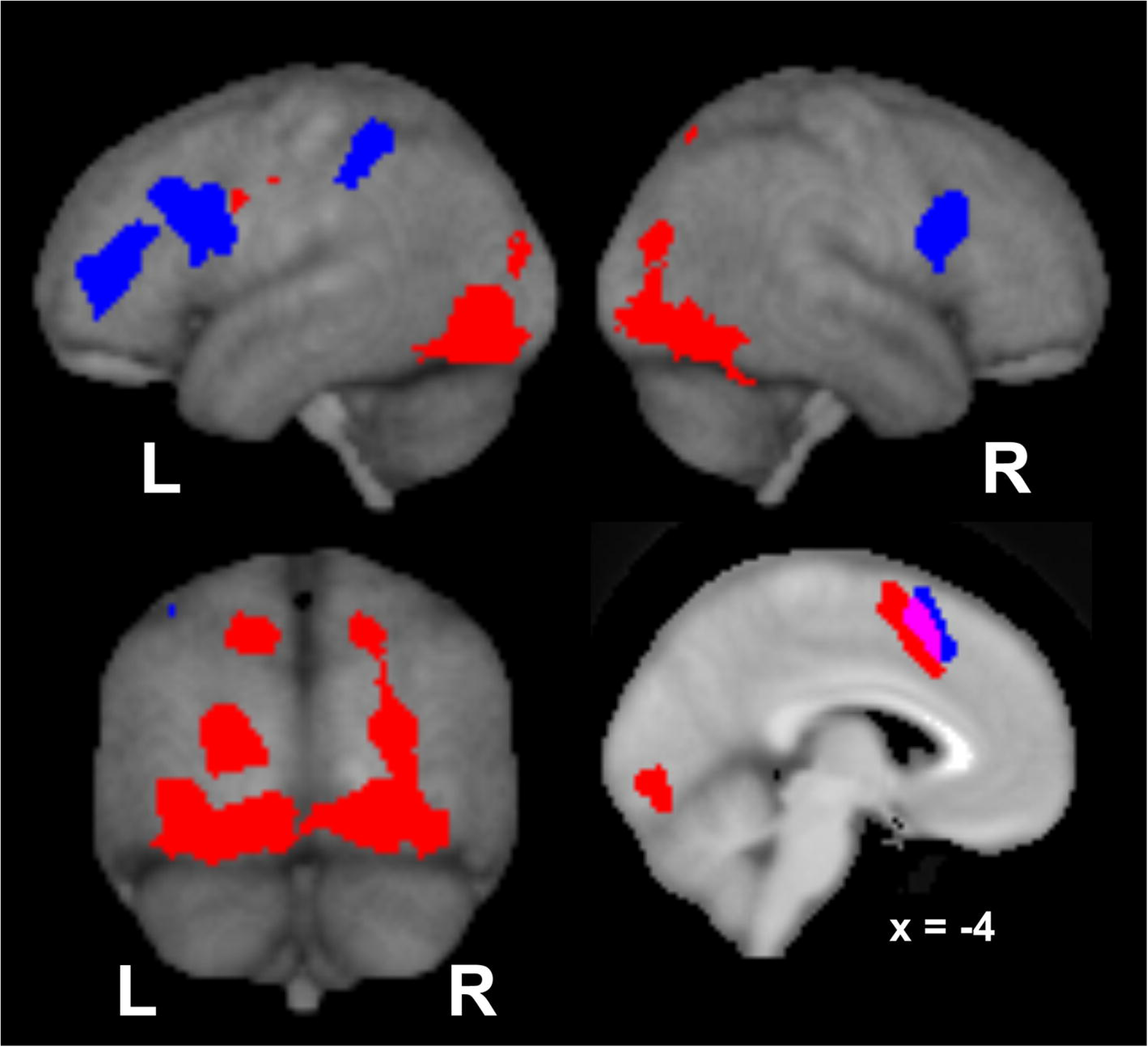
A PO and DO sentence production model based on the present findings. Red represents semantic encoding-related areas (Figure 4) and blue represents grammatical encoding-related areas (Figure 5). Overlap in the pre-SMA is indicated by pink. The activation is rendered on the skull-stripped ICBM152 MNI template pre-installed in the SPM package.

### 4.2. Neural substrates of L2 sentence production more prominent in DO than PO

To depict the distinctive neural substrates, we compared the neural activation associated with DO and PO production. As a result, activation in the pre-SMA, bilateral IFG, and left inferior parietal lobule was found, all of which are known to involve the lexical processes. As these regions did not overlap with activation of semantic encoding-related areas, except in the pre-SMA, they are likely related to the linguistic formulation process.

#### 4.2.1. IFG

According to an L1 study, the left inferior frontal regions can be dissociated into areas responsible for core syntactic computation and non-syntactic verbal working memory, with the former being located in the left pars opercularis and the latter in the left inferior frontal sulcus (Makuuchi et al., 2009). Previous studies report that increased syntactic computation demands are represented in inferior frontal and premotor areas. A lexical priming effect of the verb within a sentence aids sentence comprehension, as verb repetition shows less activation in the IFG than does the no repetition and noun repetition conditions in the posterior portion of the IFG (BA 44) and the adjacent precentral gyrus, reporting that a possible function of this region is to build syntactic representations of a sentence and determine how sentence constituents are related to each other. (Newman et al., 2009). This previous study indicates that a possible function of this region is to build syntactic representations of a sentence and determine how sentence constituents are related to each other. The increased syntactic computation involved in the reanalysis and reconstruction of sentences resulted in activation of the left IFG (BA 44/45), precentral gyrus (BA 6), and posterior temporal areas (Christensen, 2010). These previous findings are in line with our speculation that the longer RTs for DO production may be the result of PO to DO conversion, which requires retention and reordering of the phrases and tapping into syntactic working memory (Makuuchi et al., 2009).

We interpreted that activation in the right IFG does not reflect grammatical processing, as no suprathreshold activation in the right IFG was observed when a conjunction analysis of the nD > nP contrast and the “sentence production related areas (Figure 4)” was conducted. In one study, activation of the right IFG was observed when participants switched from passive to active or active to passive sentences (syntactic perturbation), and the authors discuss its role in action inhibition when subjects planned to produce a sentence from a different sentence construction (Matchin and Hickok, 2016).

#### 4.2.2. The supramarginal gyrus

The left supramarginal gyrus is involved in phonological memory, whereas the subvocal rehearsal system is associated with Broca’s area (BA 44) (Paulesu et al., 1993). Bilateral posterior parietal areas are involved in coding order information (Marshuetz et al., 2000). An L2 study that investigated neural repetition suppression using a sentence comprehension task reported German–English late-acquisition bilinguals show repetition suppression in the left MTG, the left IFG (BA 44/45), and the left precentral gyrus (BA 6) following the repetition of syntactic structure from L1 to L2 and within L2 (Weber and Indefrey, 2009). This suggests that these regions are responsible for grammatical processing, both in L1 and L2. As the critical difference between the PO and DO structure is word order, these areas may be responsible for the positional processing of the constituents of the PO and DO structures.

### 4.3. Behavioral findings

#### 4.3.1. The difference between PO and DO

Both error rate and RT were greater for DO than for PO production. This suggests that additional cognitive load is required for DO production compared to PO production. Stated more simply, DO is more difficult to produce than PO for Japanese English learners, which may explain the strong preference for PO over DO (Morishita, 2011; Nakagawa et al., 2013). One previous study showed that when Japanese English learners (N = 462) were asked to write a sentence that described a cartoon with PO or DO sentences as quickly as possible, they exhibited a clear tendency to use PO rather than DO structures (Morishita et al., 2011). This indicates that constructing DO requires greater effort than PO, which leads to PO preference.

#### 4.3.2. Cross-structural priming effect

We investigated the cross-structural (across DO and PO) priming effect. Specifically, we were interested in the priming effect by the repetition of syntactic structures (pP, dP, dD, pD) compared to that of syntax following non-syntactic structures (nP, nD).

There was a lower error rate for PO or DO sentences produced immediately after syntactic structures than after non-syntactic structures. Importantly, this effect was observed not only by repetition of identical structures (pP or dD) but also by sequential production of different structures (pD or dP). This finding indicates that PO and DO share a common process that is not shared with N trials. We experimentally eliminated the semantic encoding and grammatical encoding in N trials. Furthermore, the phonological encoding is shared across N, P, and D conditions, the effect of phonological encoding on the cross-structural priming effect is likely small. Similarly, utterance is almost identical across D, P, and N, thus the observed cross-structural priming is difficult to be explained by the utterance per se. Finally, neuroimaging results suggest the grammatical process differs between PO and DO. Thus, according to the speech production model of Bock and Levelt (1994, Figure 1), the common process is likely the semantic encoding.

The cross-structural priming effect was not clearly observed in RT measurement. Instead, the structure-specific priming effect was observed, more prominent in PO than DO. This may be caused by the sequential presentation of the cues that prompt different processes. In the present experimental task, selection cue of syntactic or non-syntactic structure comes first: black-and-white picture presentation with partial sentence or non-sentence for 2500 ms. During this period, competitive suppression between PO or DO and N may occur: Based on the competitive account (Segaert et al., 2011, 2014, 2016), a competition occurs between two structural alternatives when asked to select only one of them. Priming one candidate inhibits the other in proportion to its own likelihood to be selected. When speakers repeatedly produce syntactic structures instead of simply naming words (i.e., non-syntactic structure), the likelihood of constructing a syntactic structure compared to non-syntactic structure increases, while the production of non-syntactic structures is inhibited. Note should be made that the first cue of syntactic structure include the partial sentence, thus priming the semantic encoding process observed as the shortening of RT and decrease in error rate of both PO and DO processes. On the other hand, the second cue indicating a specific structure (PO or DO) by the colored picture follows the first cue (See Figure 2). As we measured the RT from the presentation of the colored picture to the onset of the utterance, the priming effect on the RT reflects the PO / DO selection in addition to the priming of the semantic encoding. Overall, these behavioral data support the speech production model (Figure 1), and further indicating that the difference between PO and DO processing resides in the grammatical encoding.

### 4.4. Limitations

First, we did not collect data from L1 English speakers in the present study. This is an important point, since a L1 vs. L2 comparison in the field of learning would reveal how processing and production change with higher proficiency. In many cases, speaking an L2 is not as easy as an L1. The present findings suggest that the difficulty of L2 grammatical processing is derived from grammatical encoding. To verify this, L1 speakers’ data on the same experimental tasks should be analyzed. Second, cross-linguistic studies are needed to extend our findings to English learners who speak other first languages. Analogous to the shift in the bilingual lexicon that occurs where the lexical concept is initially accessible only through L1 and eventually becomes directly accessible from L2 (French and Jacquet, 2004), L2 learners may shift from PO-biased processing to a more balanced processing as they gain proficiency. The reason for the asymmetry between PO and DO in Japanese English learners is unclear. This could simply be due to the lack of complete knowledge of DO processing (McDonough, 2006) or the cross-linguistic influence from Japanese, i.e., L1 transfer (Tokowicz and MacWhinney, 2005; Xue et al., 2013; Vaughan-Evans et al., 2014). For example, in Japanese, particles are used to mark the recipient, which is similar to marking the recipient with a preposition. The asymmetry may also be caused by greater exposure to PO than to DO, as exposure to L2 affects the preference for a syntactic structure when parsing sentences (Dussias and Sagarra, 2007). Korean English learners may show a similar pattern since they show stronger PO than DO priming effects (Shin and Christianson, 2009), which indicates a PO preference, similar to Japanese. The opposite preference has also been noted; for example, there is a preference for DO in native German speakers (Chang et al., 2015). In fluent German-English bilinguals, the production of German dative sentences primes the subsequent use of English datives and vice versa; this between-language priming is clear for DO but weak for PO, possibly due to the grammatical restrictions in German language (L1) (Loebell and Bock, 2003). Our proposed model would be strengthened if German English learners showed an opposite pattern to Japanese English learners. Third, we admit that the two syntactic structures are different in whether it emphasize (focus) on the object to be transferred or the receiver of the action, possibly confounding some of the fMRI differences between the two conditions. In order to make a distinction in such linguistic difference, modifications from the present experimental task is needed. However note that L2 learners, particularly those with low proficiency, are not necessarily aware of the difference between the two structures like native speakers of English, and thus we do not know if such confounding exists. Finally, we were not able to find any proficiency dependencies in the present study, although we collected data from various participants. There were three high-proficiency participants who were classified as CEFR C1 levels or above in our data. We performed data analysis omitting these three participants but the results did not change. Thus we concluded that the findings in the present study is proficiency independent. However, this point needs further investigation as there were only small number of high proficiency participants.

## 5. Conclusions

The present study investigated the neural basis of sentence production in L2, focusing on why DO is more difficult than PO for Japanese English learners. In sum, our findings suggest that L2 learners follow similar processing steps to L1 speakers when producing sentences. In particular, we observed distinct neural substrates underlying prelinguistic (semantic encoding) and linguistic (grammatical encoding) process. L2 semantic encoding is represented in fronto-parietal-occipital regions, while grammatical encoding is represented in the fronto-parietal regions. We conclude that one of the reasons why L2 speaking is challenging is because additional computation is required for grammatical encoding, conducted mainly in the left inferior frontal regions.

## Author Contributions

EN: Conceptualization, Methodology, Validation, Data collection & analysis, Writing & Editing, Visualization, Funding acquisition. TK: Conceptualization, Methodology, Data collection & analysis, Editing, Supervision. MS: Validation, Data collection & analysis. KS & KM: Validation. HaY: Conceptualization. HiY: Conceptualization, Editing, Funding acquisition. NS: Conceptualization, Writing & Editing, Supervision, Project administration, Funding acquisition. All authors reviewed the paper. All authors contributed to the article and approved the submitted version.

## Footnotes

1 An asterisk (*) indicates grammatically incorrect text.

